# Interpersonal Synchronization in Mother-Child Dyads: Neural and Motor Coupling as a Mechanism for Motor Learning and Development in Preschoolers

**DOI:** 10.1101/2024.10.03.616469

**Authors:** Péter Nagy, Luca Béres, Brigitta Tóth, István Winkler, Betty Barthel, Gábor P. Háden

**Affiliations:** Institute of Cognitive Neuroscience and Psychology, HUN-REN Research Centre for Natural Sciences, 1117 Budapest, Magyar tudósok körútja 2., Hungary; Department of Artificial Intelligence and Systems Engineering, Faculty of Electrical Engineering and Informatics, Budapest University of Technology and Economics, 1117 Budapest, Magyar tudósok körútja 2., Hungary; Department of Cognitive Science, Faculty of Natural Sciences, Budapest University of Technology and Economics, 1111 Budapest, Egry József utca 1., Hungary; Institute for Special Education of Atypical Behavior and Cognition, Bárczi Gusztáv Faculty of Special Needs Education, Eötvös Loránd University. 1097 Budapest, Ecseri út 3., Hungary; Department of Telecommunication and Artificial Intelligence, Faculty of Electrical Engineering and Informatics, Budapest University of Technology and Economics, 1117 Budapest, Magyar tudósok körútja 2., Hungary

## Abstract

Interpersonal movement synchrony (IMS) and brain-to-brain coupling play a crucial role in social behavior across species. In humans, IMS is often studied in structured tasks that require specific body movements, while spontaneous, unstructured movements have received less attention. In this study, we investigated both structured and spontaneous motor coordination in mother-child dyads. We recorded upper-body kinematics and dual-EEG from mothers and their preschool children during motor tasks and spontaneous face-to-face interactions. Our findings show that mother-child dyads synchronize their movements and neural activity, particularly in gamma band oscillations. This motor and neural synchrony evolves across task repetitions, with a strong correlation between motor and neural measures. Further, we observed that only motor synchronization was significantly related to the child’s motor development stage, as assessed by the Movement Assessment Battery for Children. These results suggest that gamma band brain-to-brain coupling reflects joint motor coordination and mutual adaptation shaped by structured tasks and spontaneous interpersonal interactions.

## Introduction

Joint action, the coordination of actions between individuals, is crucial for human success and has gained significant research attention recently (Sebanz et al., 2006). Synchrony, defined as acting on a shared time reference frame, is essential to all forms of human interaction, from turn-taking communication to joint action towards shared goals, all towards larger-scale social interaction. Mothers and children mutually synchronize from birth and throughout development (Jaffe et al., 2001), and children benefit from the scaffolding synchrony, which provides better predictions about the surrounding world (i.e., learning). Tracking the motion of multiple agents is an easily accessible way of assessing synchrony. With the recent advances in neuroscience, we now have unprecedented access to synchrony on a neural level. Utilizing this access, we may answer questions such as how preschool children and caregivers synchronize when performing and learning motor tasks, how synchrony changes with practice, and how this is achieved on a behavioral and neural level.

By testing preschool children performing motor synchronization tasks with their mothers, we try to gain insights into the development of synchronization and the factors affecting task performance, including the tightness of synchronization, the rigidity of the task, and the overall motor performance of the child. We assume that further insights may be gained by measuring both behavior and neural data that can be analyzed in the same time frame.

### Neural brain-to-brain synchronization during interpersonal situations and its developmental aspects

Being ‘in sync’ or ‘on the same wavelength’ with someone are phrases often used to describe the feeling of a unique bond or closeness with someone. Humans are exceptionally good at attuning to one another in the physiological and behavioral sense. Research in the past decade has shown that this attunement is substantiated in the brains of interacting individuals, which researchers often refer to as interpersonal neural synchronization or brain-to-brain synchronization. Recent advances in social neuroscience, e.g., the simultaneous measurement of two or more interacting individuals - frequently termed ‘hyperscanning’ (Babiloni and Astolfi, 2014) - have allowed researchers to study synchronization in situations with higher ecological validity. As a result, it has been demonstrated that synchronization is prevalent across a range of different contexts, e.g., during collaborative problem-solving (Nguyen et al., 2020), everyday conversations (Kinreich et al., 2017), playing a musical duet with someone (Mueller et al., 2013), etc.

Studies have explored various aspects of joint action, including joint attention, action observation, and task sharing (Sebanz et al., 2006). The cognitive perspective emphasizes joint coding and mirror neuron theories (Rizzolatti & Craighero, 2004; Rizzolatti, Cattaneo, Fabbri-Destro & Rozzi, 2014), while the behavioral dynamics approach focuses on emergent patterns like synchronization (Schmidt et al., 2011). Neuroimaging research has revealed that joint action contexts modulate motor-related brain activity, with dissociable activity for one’s actions, partner’s actions, and the joint action itself (Bolt & Loehr, 2020). Additionally, between-brain coupling of motor-related oscillatory activity has been observed (Bolt & Loehr, 2020). Joint actions involving complementary actions or leader-follower roles elicit unique motor activity (Bolt & Loehr, 2020). Mental representations, shared sensorimotor information, and general coordination mechanisms contribute to successful joint action (Vesper et al., 2017; Keller et al., 2014).

Brain-to-brain synchrony is primarily supported by specific brain oscillations and regions involved in social cognition and communication. Key brain regions supporting this synchrony include the prefrontal cortex, which is involved in higher-order cognitive functions such as decision-making and empathy, and the temporoparietal junction (TPJ), which plays a crucial role in perspective-taking and understanding others’ intentions (Martin et al., 2020). The superior temporal sulcus (STS) is also important for processing social cues, like facial expressions and gaze direction (Paracampo et al., 2018). These brain regions and synchronized oscillatory activity create a neural foundation for effective social communication and cooperation.

Brain oscillations facilitate communication between brain regions and support inter-brain synchrony during social interactions. Lower frequency oscillations (<25 Hz) provide a temporal framework for information carried by higher frequency activity (>40 Hz), enabling effective communication between brain areas. During social interactions, interbrain synchronization emerges in the alpha-mu band between the right centroparietal regions, which act as crucial functional hubs (Dumas et al., 2012). In naturalistic social interactions, brain-to-brain synchrony occurs in gamma rhythms localized to temporal-parietal structures, particularly among romantic couples. This neural synchrony is linked to behavioral synchrony, social gaze, and positive affect, highlighting the role of social connectedness in two-brain coordination (Kinreich et al., 2017).

Numerous studies have focused on brain-to-brain synchrony in adults, and there is increasing interest in exploring the phenomenon in parent-child dyads. Interpersonal synchrony is present from birth and shapes social interactions between individuals throughout their lives. Rhythmic communication often achieves synchrony (Hoehl et al., 2021). Research on joint action and motor coordination in older children reveals important developmental milestones. Around age 7, children begin synchronizing actions with partners during joint tasks, showing improved reaction times and reduced variability (Satta et al., 2017). By age 8, solo performance accuracy peaks, and children implement online monitoring of peers during joint actions (Satta et al., 2017). Children with autism spectrum disorder (ASD) struggle with joint action coordination when relying solely on kinematic information, displaying less accurate and slower movements compared to typically developing (TD) children (Fulceri et al., 2018). In real-world collaborative tasks, children with ASD are less likely to benefit from peer collaboration and show reduced step synchronization, though these differences diminish with adult scaffolding (Trevisan et al., 2021). Movement coordination characteristics in parent-child dyads influence goal attainment and relate to children’s first-order theory of mind but not second-order theory of mind (Białek et al., 2022).

Research on brain-to-brain synchrony reveals its importance in child development and parent-child interactions. Neural synchrony, involving the coordination of oscillatory activity, plays a crucial role in cortical network development from childhood through adolescence (Uhlhaas et al., 2009;). Parent-child brain synchrony, particularly in prefrontal regions, has been observed during cooperative tasks and is associated with improved performance and emotion regulation (Reindl et al., 2018). This synchrony is specific to parent-child dyads and is not observed with strangers or during competitive interactions. Recent studies have linked parent-child brain synchrony to various factors, including interaction type, social cues, child characteristics, and environmental influences (Alonso et al., 2024). The development of neural synchrony continues into early adulthood, coinciding with changes in myelination and GABAergic neurotransmission (Uhlhaas et al., 2010). Understanding brain-to-brain synchrony may provide insights into typical development and neurodevelopmental disorders.

### Interpersonal movement synchrony and its development

Similarly to dual EEG, motion-tracking systems capable of recording multiple agents at comparable speeds to EEG were used to track adult and infant interactions (Cuadros et al., 2019), and various algorithms can analyze both neural and motion data (e.g. Issartel et al., 2015, Bastos, & Schoffelen, 2016). Interpersonal motor synchronization (IMS) involves coordinated movements between individuals, which can have significant social implications. Even at a very early age, there is a balance in mother-infant dyads between autonomy and synchrony (Beebe et al., 1982). Similarly, when two agents move together, there is a balance between coupling and autonomy (Dumas et al., 2014). Research has shown that IMS increases prosocial behavior in infants, with 14-month-olds exhibiting more altruistic actions after synchronous bouncing with an experimenter (Cirelli et al., 2014). The dynamics of IMS can be observed in various contexts, from structured tasks like hand-clapping to more complex interactions such as conversations. Importantly, interpersonal perception plays a crucial role in joint actions, with negative relationships improving performance in guided interactions but hindering free interactions that require mutual adjustments (Sacheli et al., 2012).

All motor learning theories agree that any motor task improves with repeated performance (Wolpert et al., 2001). However, even repetition requires the ability to represent the goal of the task and the necessary movements to achieve said goal. One of the key challenges in development is mastering one’s body’s ability to perform movements varying in complexity, from raising one’s head to performing a coordinated dance routine with several other people. As the human body has a high degree of freedom, infants and young children spend years mastering coordinated action. One of the ways this process is hypothesized to occur is through representing the perceived body motions of others in the so-called mirror system (Cuevas et al., 2014). The mirror system, as well as other sources of information (visual self, proprioception), are then used in creating body schemes that represent different body parts and the associated actions (Brownell et al., 2010) that can be performed with these body parts. These schemes are flexible and are extended when learning new motor skills, such as running or using an object (Cardinali et al., 2012). The schemes are also helpful in learning from others through imitation or direct teaching (Meltzoff & Marshall, 2020; 2018).

Motor synchronization plays a crucial role in motor learning and performance. During the acquisition of complex bimanual tasks, cortico-spinal synchronization in the beta band correlates with improved performance (Houweling et al., 2010). As learning progresses, global decreases in brain activity are observed, with specific increases in the left primary motor cortex (M1) and right cerebellar regions associated with improved synchronization (Steele & Penhune, 2010). Inter-hemispheric gamma synchronization between motor areas decreases, possibly reflecting reduced attentional demands (Houweling et al., 2008). Motor unit synchronization, which exhibits acute and chronic plasticity, enhances force development during rapid contractions and coordinates muscle synergies (Semmler, 2002). Additionally, alpha modulation in the cerebellum and increased bilateral M1 coupling around movement frequency indicate improved motor timing (Houweling et al., 2008). These findings highlight the importance of neural synchronization in various brain regions for motor learning and performance optimization.

### Aims and hypothesis

There is growing evidence that interpersonal synchronization - acting on a shared time reference frame with others - is prevalent across various contexts. Still, its effects on social behavior are only beginning to be explored. Many studies have focused on brain-to-brain synchrony in adults, and only a few have investigated the phenomenon in parent-child dyads, with fewer still concentrating on motor learning. In the present study, we tested parent-child dyads through two tasks requiring different forms of synchrony and having various degrees of freedom. The tasks will be repeated twice to assess motor learning and how it benefits from synchrony between the mother and child. The tasks were selected to be challenging but feasible for the participants to avoid a ceiling effect on learning. The primary objective of this study is to examine the relationship between motor coordination and brain-to-brain synchrony in mother-child dyads during joint motor tasks. The study uses dual-EEG scanning and upper-body motion tracking to investigate how motor and neural synchronization evolves across task repetition.

Additionally, the study explores how much a child’s motor development, as assessed by the Movement Assessment Battery for Children, influences motor and neural synchrony within the dyads. We hypothesize that greater motor-neural synchronization will correlate with more advanced motor task coordination execution, while reduced synchrony may serve as an early indicator of developmental delays (motor skills or joint attention).

## Materials and Methods

### Participants

Twenty-four mother-child dyads (N=24) participated in the study. Due to technical issues during recordings, data from four mother-child dyads had to be excluded from the analysis. All mother and children participants were right-handed, and the children were evenly split by gender (10 girls and 10 boys). The average age of the mothers was 40.5 years (SD = 5.4), while the children had a mean age of 71.75 months (SD = 7.5). On average, the mothers reported 7.3 hours of sleep per night, and the children averaged 9.125 hours of sleep. None of the participants had any history of neurological or psychiatric disorders.

Dyads were recruited through a database of families who already participated in previous experiments and consented to be contacted. All mothers provided informed consent to the experiment, and the study was conducted in full compliance with the Declaration of Helsinki and relevant national regulations. The United Ethical Review Committee for Research in Psychology (EPKEB), Hungary, granted ethical approval. Participants received no compensation except for a small gift for the children at the end of the experiment.

Before moving on with the experimental procedures, all children completed the Movement Assessment Battery for Children, Second Edition (MABC-2; Henderson, Sugden & Barnett, 2007; Smits-Engelsman et al., 2008), a standardized test to assess motor skills in children aged 3-16. The MABC-2 evaluates three domains: manual dexterity, aiming and catching, and balance. It is commonly used to identify motor difficulties, with the total test score providing insight into the severity of any movement problems. The test categorizes results into three zones: the red zone, indicating significant movement difficulties; the amber zone, signaling that the child is "at risk" and may require monitoring; and the green zone, where no movement difficulties are detected. None of the children in this study showed signs of movement difficulties. Their average Total Test Score was 84.80 (SD = 9.21), and the average Standard Score was 11.45 (SD = 2.31). Their average Percentile Rank was 65.70 (SD = 24.14(, comfortably in the green zone, above the 15th percentile, supporting the conclusion that no child required monitoring for motor difficulties based on the assessment.

### Stimuli and recording setup

The laboratory for the experiments was set up in a 5-meter by 5-meter room with artificial lighting. An adjacent room separated by a 2-way mirror was used to control the experiment and data collection. In the center of the room, two adjustable-height stools were set up parallel to the 2-way mirror, with an additional step-up stool for children that was set to allow for the firmly putting of both feet on the ground (stool). The mother always sat to the right when viewed from the control room. 12 Naturalpoint (Naturalpoint Inc., Corvallis, OR, USA) OptiTrack Primex13 cameras were arranged around the room for motion capture. Motion tracking data was recorded with Motive (version 2.3) software at 240 frames per second (fps). Reflective markers were attached to both participants’ upper bodies and heads using the OptiTrack Baseline Upper (25) marker layout. EEG was recorded using two 64-channel ActiChamp+ EEG amplifiers connected to 64-channel R-Net electrode systems (RNP-AP-64 standard layout and channel assignment; Brain Products GmbH, Gilching, Germany) with a sampling rate of 1 kHz. The audio was recorded at a 48 kHz sampling rate using a Focusrite (Focusrite Ltd. High Wycombe, UK) Scarlet 4i4 USB audio interface. Color video (720p, 60 fps) was recorded using a Sony HDR-CX550 camera (Sony Corp., Tokyo, Japan) and a USB video capture device behind the 2-way mirror. All data streams (EEG1, EEG2, Motion tracking, Audio, Video frame count) were recorded (in addition to local recording for all streams except EEG1 and EEG2) through the Lab Streaming Layer (Kothe, 2014) using LabRecorder (version 1.16.4), LSL connector for the actiCHamp (Brain Products GmbH, 2022) for the EEG streams and custom in-house software for the other streams. A custom-built clapperboard with attached reflective markers for motion tracking, capable of producing sound, light, and sending a TTL trigger toward the EEG, was used to provide additional synchronization cues.

Stimuli were presented using Matlab (MathWorks Inc., Natick, MA, USA) and Psychtoolbox (Kleiner, Brainard, and Pelli, 2007). The auditory stimulus to signal the beginning of the tasks was a 5 ms long 1500 Hz pure-tone beep, presented through speakers connected to a Maya22 USB sound interface (ESI Audiotechnik GmbH, Leonberg, Germany). Visual stimuli were presented on a 26” monitor behind the child, only visible to the mother. A visual stimulus signaling the beginning of tasks was a uniform green display. A second stimulus was a two-by-four grid of eight different colors filling up the entire screen, and the order of the colors in the grid was randomized for each repetition. A custom-built 0.6-meter by 0.78-meter plywood labyrinth built on a thin plywood base, with eight color-coded target locations with a small rubber ball fitting the labyrinth’s paths, was used for one of the tasks.

### Procedure

The motion tracking system was calibrated before each experimental session. Upon arrival, participants were familiarized with the laboratory, and the child was administered the MABC-2 test in a separate testing room while the mother filled out a questionnaire of basic demographic data and questions regarding the motor and social development of the child. Upon finishing, both participants were fitted with reflective markers and EEG electrodes. Once motion tracking and EEG recordings were confirmed to work, participants took part in 3 games with a fixed order and repeated the sequence once (Figure 1), for the current study, data from the Mirror Game and the Labyrinth Game is analyzed. Participants could take rests between games when requested and were explicitly prompted to rest after the completion of the first sequence.

**Figure 1.**
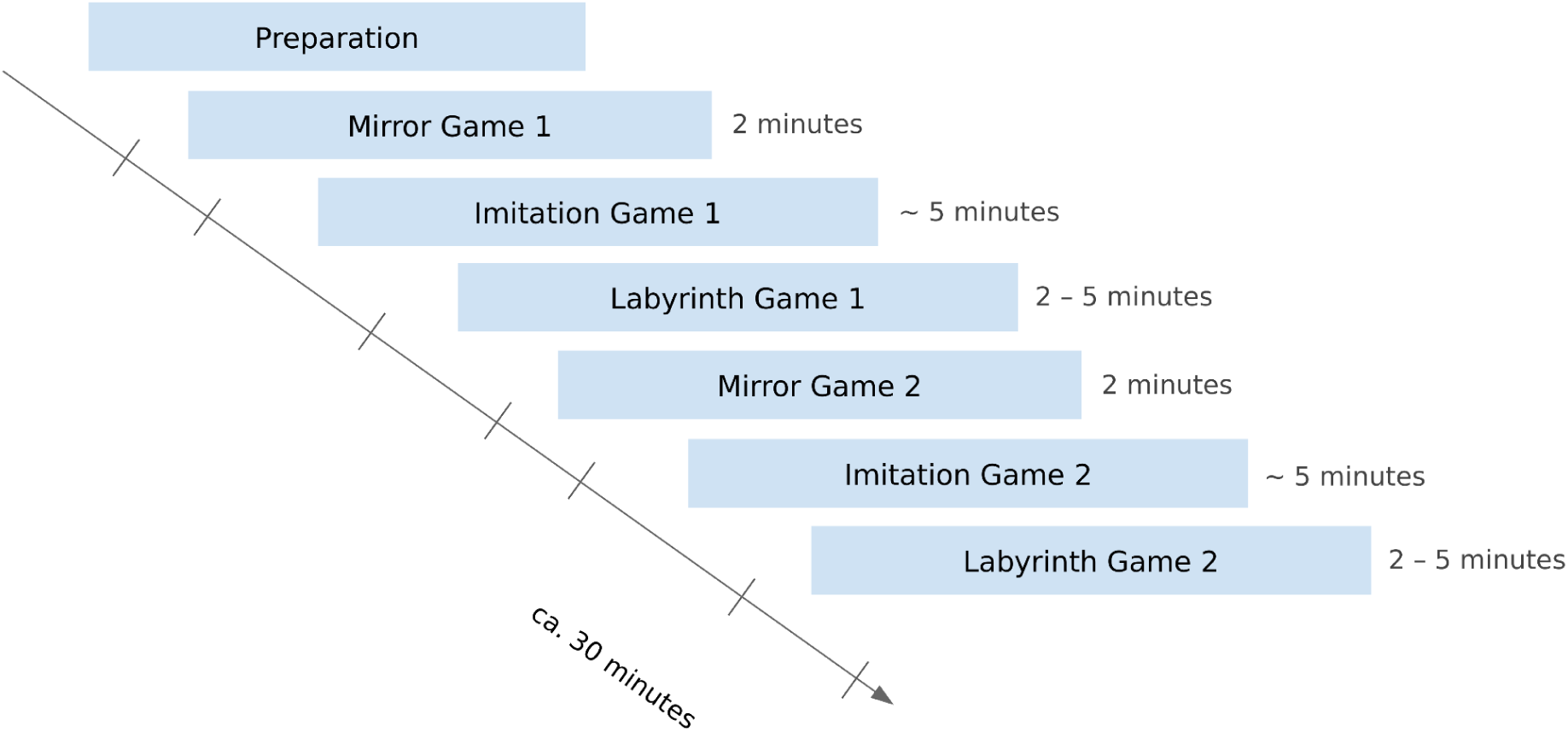
Timeline of experimental tasks

The Mirror Game is based on the paradigm of Noy et al. (2011). Participants sat with their hands extended in front of their bodies, their palms almost touching. The mother was asked to improvise movements with her, and the child was instructed to follow (“mirror”) these movements closely. The game lasted for 2 minutes and started with an audiovisual prompt.

In the Labyrinth Game (Figure 2), participants hold a labyrinth board with eight color marked positions. Their task was to move a rubber ball in the labyrinth to the marked positions in a set order by tilting the labyrinth board together. The mother saw the order of the color-coded positions on a screen behind the child.

**Figure 2.**
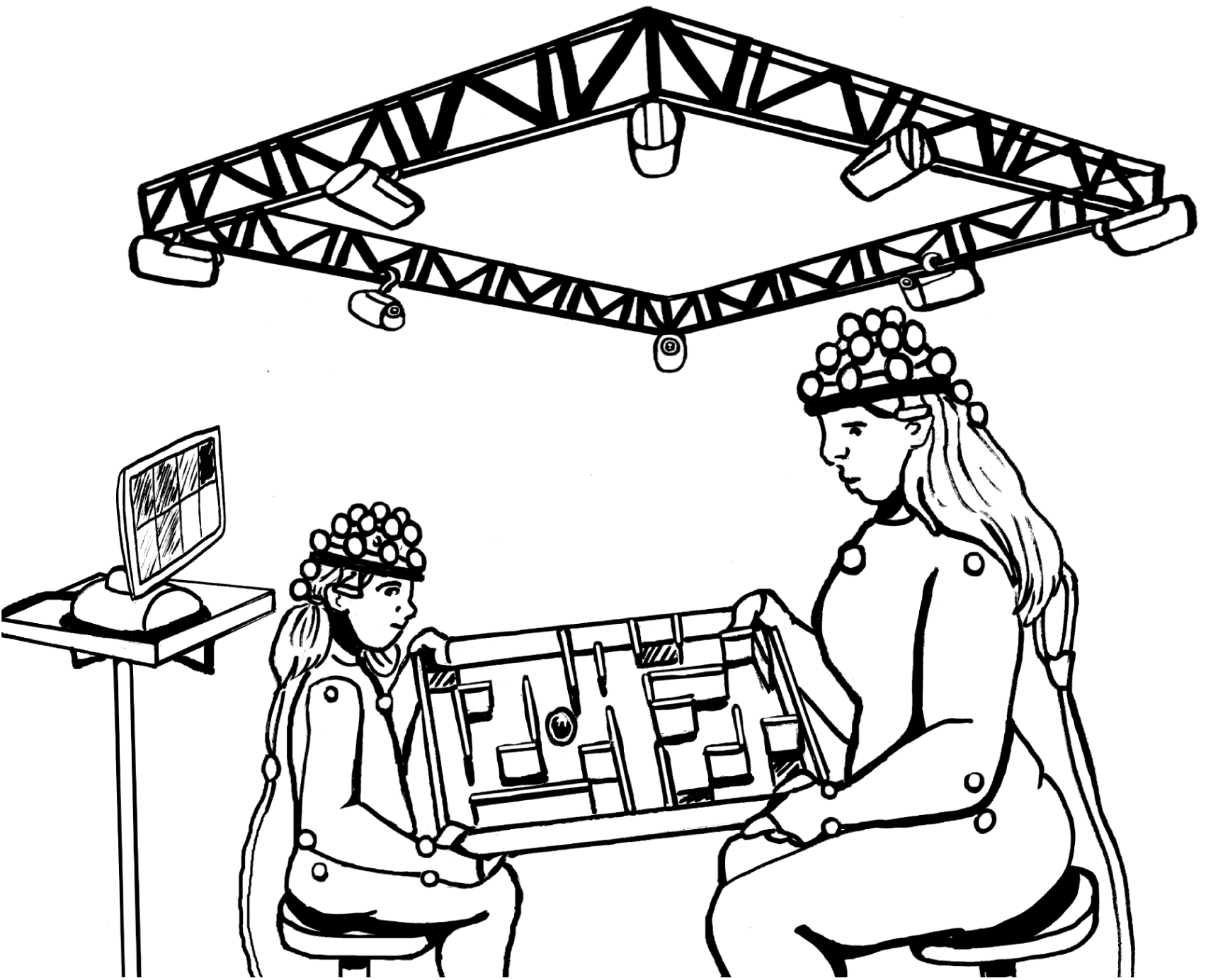
Schematic representation of the setup for the Labyrinth Game. The mother and child sit facing each other at a distance that enables both to hold the labyrinth table comfortably at the side near the corner. The screen behind the child shows the order of the color-coded target positions.

### EEG Preprocessing

Preprocessing of EEG was done using MATLAB (2024a; MathWorks) and EEGLAB toolbox (v2024.0; Delorme and Makeig, 2004). First, the EEG signals were re-referenced to the common average and bandpass filtered between 0.5 Hz and 80 Hz (Hamming windowed sinc FIR filter, a cascade of a high pass and a low pass filter), with notch filtering at around 50 Hz (47.0 - 53.0 Hz Hamming bandstop filter). A maximum of six wrong EEG channels per participant were interpolated using the spherical interpolation method implemented in EEGLAB. The Infomax algorithm of Independent Component Analysis (ICA) implemented in EEGLAB was employed for artifact removal, with decomposition to a principal component subspace; 32 principal components were retained. ICA components constituting artifacts related to blinks and horizontal eye movements were removed via visual inspection based on the components’ topographical distribution, temporal activity, and frequency content. Maximum 3 ICA components were removed. Preprocessed EEG signals were bandpass filtered into five frequency bands with zero-phase FIR filters: delta (0.5-4 Hz), theta (4-8 Hz), alpha (8-13 Hz), beta (13-30 Hz), gamma (30-80 Hz). EEG signals were used to estimate interpersonal neural synchronization by functional connectivity (Balconi and Vanutelli, 2017; Yun et al., 2012). Inter-brain connections were estimated between homologous electrode groups of the mother and the child. The following groups of electrodes were investigated: left frontal (F1, F3, F5, AF3), right frontal (F2, F4, F6, AF4), left central (FC1, FC3, FC5, C1, C3, C5), right central (FC2, FC4, FC6, C2, C4, C6), left parietal (CP1, CP3, CP5, P1, P3, P5), right parietal (CP2, CP4, CP6, P2, P4, P6). The average of signals within each group was calculated, and Amplitude Envelope Correlation (AEC) was computed between the signals of homologous groups (e.g., the signal of the left parietal electrode group of the mother and the signal of the left parietal electrode group of the child). Synchronization between the amplitude envelopes of band-limited oscillations has been initially mainly studied using MEG data (O’Neill et al., 2015). More recently, amplitude envelope-based connectivity estimation has been successfully applied to investigate resting-state FC networks based on EEG signals (Coquelet et al., 2020). In the first step of AEC estimation, continuous signals were segmented into non-overlapping epochs of length 5 s. Next, for each epoch, the envelope of the signals corresponding to the two tested electrode groups was computed as the magnitude of the analytical signal. Then, Pearson’s correlation coefficient was calculated between the envelopes. Finally, correlation values were averaged over all epochs. The epoch length of 5 s was chosen based on a previous study (Nagy et al., 2024).

### Motion Synchrony

To analyze the synchrony between the mother’s and the child’s motion, the magnitude squared wavelet coherence (Fujiwara and Daibo, 2016) of the motion energy of selected marker groups was calculated. The following markers were considered: two markers on each shoulder, one marker on each upper arm, one marker on each elbow, and two markers on each wrist—the squared velocity of the markers’ estimated motion energy. To calculate the squared velocity, the derivative of position with respect to time was computed and squared separately along the x, y, z directions. Finally, squared velocities along the x, y, z directions were summed. Magnitude squared wavelet coherence (Morlet wavelets, 12 voices per octave) of the motion energy was computed between the following groups of markers:

- Total energy of both arms of the mother and total energy of both arms of the child (will be referred to as both arms)
- Total energy of both hands of the mother and total energy of both hands of the child (will be referred to as both hands)
- The energy of the **P**arent’s **L**eft **A**rm and energy of the **C**hild’s **R**ight **A**rm (referred to as PLACRA)
- The energy of the **P**arent’s **R**ight **A**rm and energy of the **C**hild’s **L**eft **A**rm (referred to as PRACLA)
- The energy of the **P**arent’s **L**eft **H**and and the energy of the **C**hild’s **R**ight **H**and (referred to as PLHCRH)
- Energy of the **P**arent’s **R**ight **H**and and energy of the **C**hild’s **L**eft **H**and (referred to as PRHCLH)

For each marker group, the motion energy of the group was calculated as the sum of the energy of the markers in the group:

- Arm (right of left): two markers on the corresponding shoulder, one marker on the corresponding upper arm, one marker on the corresponding elbow, two markers on the corresponding wrist
- Hand (right or left): two markers on the corresponding wrist
- Both arms: two markers on the left shoulder, two markers on the right shoulder, one marker on the upper left arm, one marker on the upper right arm, one marker on the left elbow, one marker on the right elbow, two markers on the left wrist, two markers on the right wrist
- Both hands: two markers on the left wrist, two markers on the right wrist

In the mirror game, the first 90 seconds of the game were considered in the coherence analysis. In the labyrinth game, the first 120 seconds of the game were considered in the coherence analysis. Magnitude-squared wavelet coherence characterizes the correlation between signals in the time-frequency plane. In both games, coherence values were averaged over time. The cone of influence was excluded from the averaging. Two frequency bands were selected: 0.1 - 1 Hz (low-frequency band) and 1 - 4 Hz (high-frequency band). In each frequency band, coherence values were averaged over frequency. This procedure resulted in two aggregated coherence values (one for the high-frequency band, one for the low-frequency band) for each marker group of each pair in each game.

### Statistical Analysis

As a first step of the statistical analysis, neural and motion synchronization measures were selected using pseudo pairs. Pseudo pairs are random pairings of participants who did not play the games together (e.g., a dyad’s mother designated by ‘01’ with the dyad’s child selected by ‘05’). As the number of real mother-child pairs was 20, the number of pseudo pairs was 380. Only synchronization measures were included in further analyses, which showed statistically different values for real pairs compared to pseudo pairs. Two-sample t-tests compared real and pseudo pairs.

Selected neural and motion synchronization measures were included in four different types of analysis.

1. Each measure was compared between the first and second repetitions of the mirror and labyrinth games using a paired-sample t-test.
2. The correlation was computed between measures of motion synchronization and measures of neural synchronization.
3. Measures of motion synchronization and neural synchronization were correlated with the movement assessment’s Total Test Score.
4. Measures of motion synchronization and neural synchronization were correlated with the times needed to complete the first and second labyrinth games.

As an additional analysis step, the times needed to complete the first and second labyrinth games were compared by a paired-sample t-test and correlated with the movement assessment’s Total Test Score.

Pearson’s correlation coefficient estimated correlation values, which were tested to see if they significantly differed from zero using Student’s t-distribution. An alpha level of 0.05 was used in all statistical tests (Bonferroni-corrected against multiple comparisons). Effect sizes were estimated using Cohen’s *d*.

## Results

### Comparing real and pseudo pairs in motion synchronization

Motion synchronization was estimated for six marker groups (both arms, both hands, PLACRA, PRACLA, PLHCRH, PRHCLH) in the low-frequency band (0.1 - 1 Hz) and in the high-frequency band (1 - 4 Hz). In the mirror game, all six marker groups showed significantly higher coherence for real pairs than pseudo pairs in both games and frequency bands (Supplement Table A1-A4, illustration for both hands in Figure 3). In the labyrinth game, only the high-frequency band showed significantly higher coherence for real pairs compared to pseudo pairs (both arms, both hands, PLACRA, PLHCRH, PRHCLH in the first labyrinth game and both arms, both hands, PRACLA, PRHCLH in the second labyrinth game, Supplement Table A5-A8).

**Figure 3.**
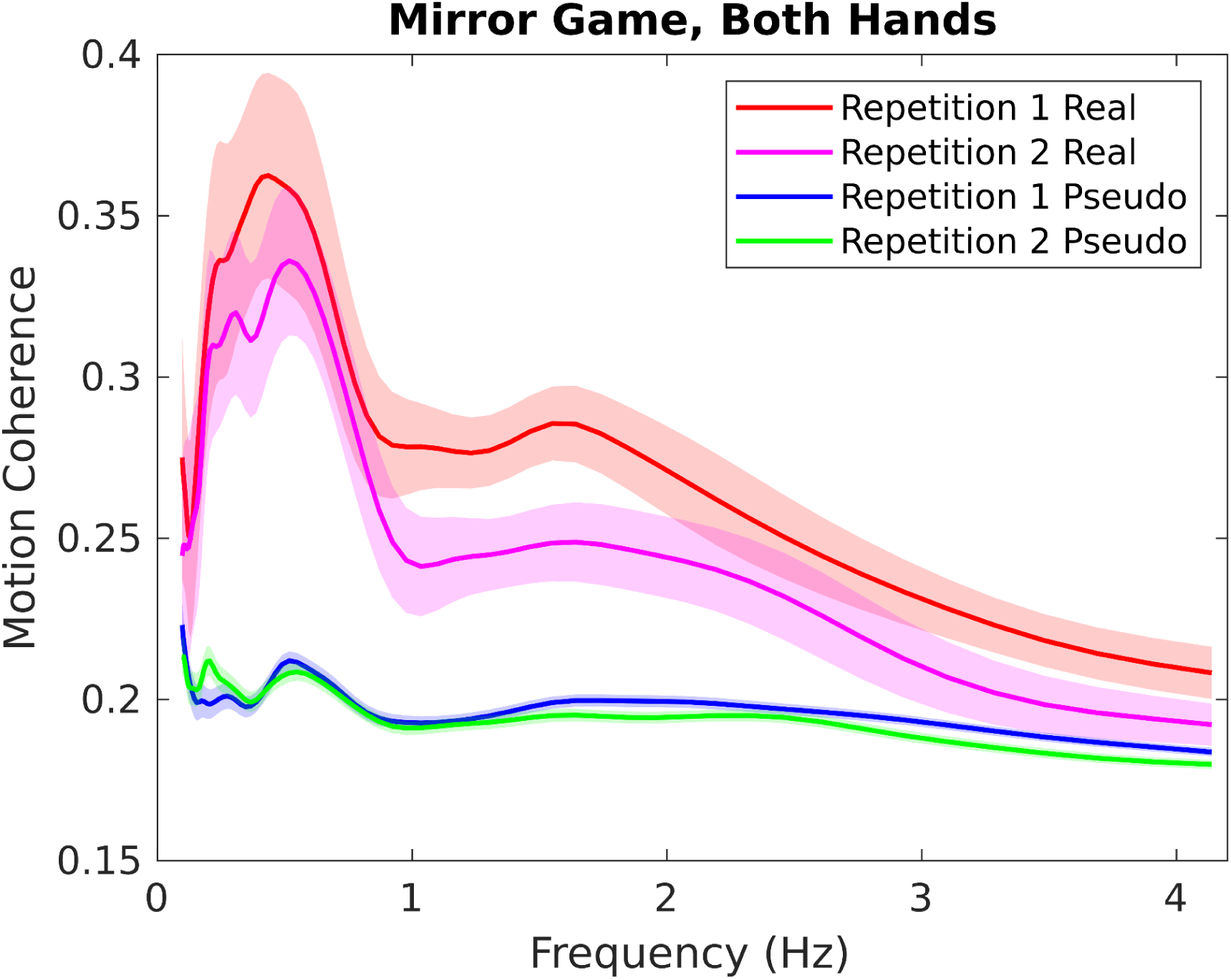
Group-average magnitude-squared wavelet coherence of real and pseudo pairs for the marker group referred to as both hands in the first and second repetition of the mirror game, in the frequency range between 0.1 - 4 Hz. Coherence values were averaged over time. A shaded area marks the standard error of the mean.

As a further illustration, we give the group-level mean value of coherence for the three marker groups producing the highest values, averaged over the two repetitions and divided by the group-level mean value of pseudo pairs. Low-frequency band in the mirror game: PRHCLH (1.8796), PLHCRH (1.7548), PRACLA (1.5356). High-frequency band in the mirror game: PLHCRH (1.3676), PRHCLH (1.3654), both hands (1.2674). Low-frequency band in the labyrinth game: both hands (1.0985), PLACRA (1.0594), PLHCRH (1.0585). High-frequency band in the labyrinth game: PLHCRH (1.1568), PRHCLH (1.1542), PLACRA (1.1436).

### Comparing real and pseudo pairs in neural synchronization

Neural synchronization was estimated by computing AEC between signals of homologous EEG electrode groups (left frontal, right frontal, left central, right central, left parietal, right parietal) of the mother and the child in the delta, theta, alpha, beta, and gamma frequency bands. Significantly different AEC values for real pairs compared to pseudo pairs were found only in the gamma band and the first games. In the first mirror game, the right frontal region (*t*(398) = 2.75, *p* = 0.0379, *d* = 0.63), the left parietal region (*t*(398) = 3.27, *p* = 0.0070, *d* = 0.75), and the right parietal region (*t*(398) = 3.14, *p* = 0.0109, *d* = 0.72) showed significantly higher AEC values for real pairs than for pseudo pairs, in the gamma band. In the first labyrinth game, the left central region (*t*(398) = 2.71, *p* = 0.0423, *d* = 0.64), the right central region (*t*(398) = 2.70, *p* = 0.0434, *d* = 0.64), and the right parietal region (*t*(398) = 4.47, *p* = 0.0001, *d* = 1.05) showed significantly higher AEC values for real pairs than for pseudo pairs, in the gamma band (Figure 4).

**Figure 4.**
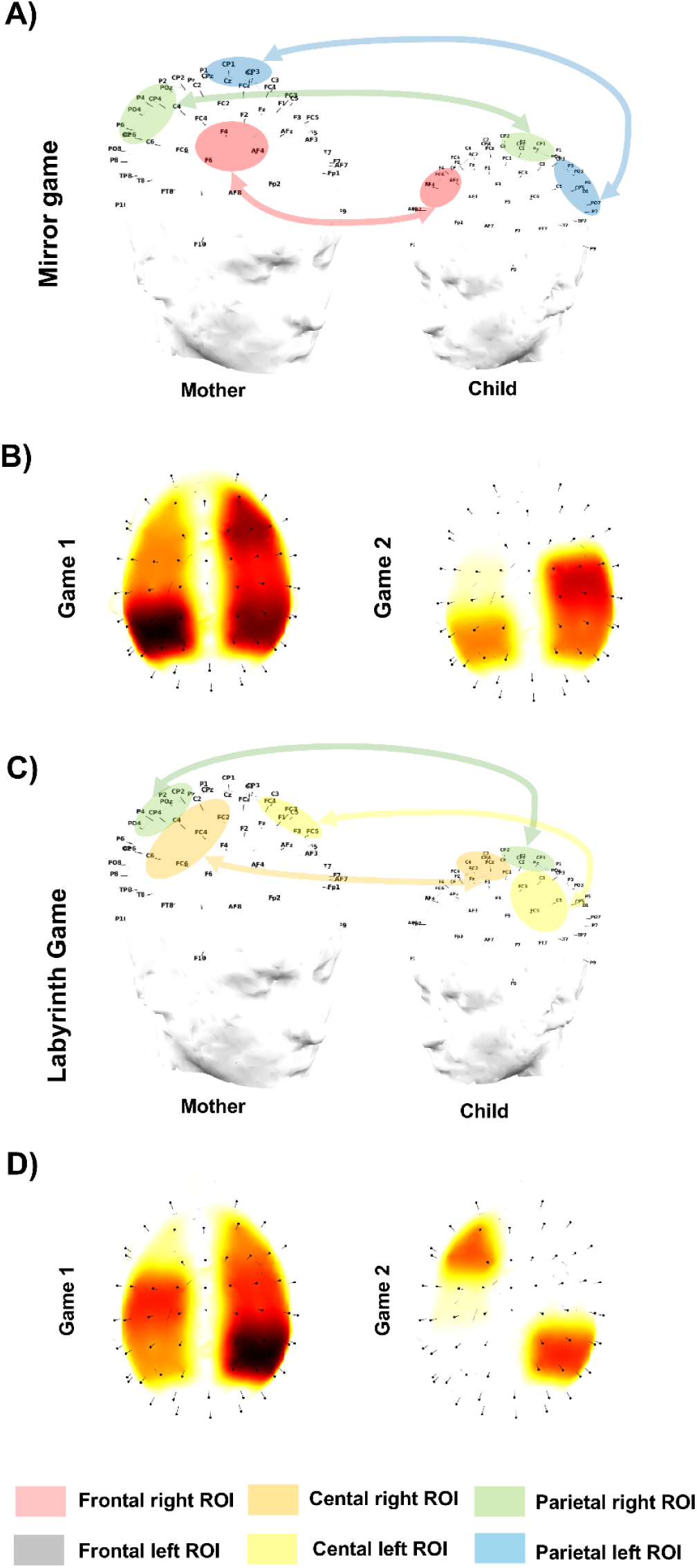
A) Significant gamma band brain-to-brain synchrony was measured in the first mirroring game. Note color indicates the ROI. B) Scalp distribution of brain-to-brain synchronization measured in mirroring game first and second attempts. Note that the synchrony evaluated for each ROI only represents the channel location as dots on the scalp. C) Significant gamma band brain-to-brain synchrony was measured in the first Labyrinth D) Scalp distribution of brain-to-brain synchronization in the Labyrinth game’s first and second attempts.

### Comparing motion and neural synchronization between two repetitions of the games

Each measure of motion synchronization and neural synchronization was compared between the first and second repetitions of the mirror game and between the first and second repetitions of the labyrinth game. Most measures showed smaller values for the second repetition than the first one. Still, a statistically significant difference was found only in the motion coherence in the high-frequency range for the marker group referred to as both hands (*t*(19) = 3.40, *p* = 0.0182, *d* = 0.64).

### Correlating motion synchronization with neural synchronization

Pearson’s correlation coefficient was calculated between measures of motion synchronization and the corresponding measures of neural synchronization. Significant correlations were found only in the first mirror game and only for the left parietal region (gamma band). In the first mirror game, a significant correlation was found between motion coherence for both hands (low-frequency) and AEC for the left parietal region (*r* = 0.77, *p* = 0.0013); between motion coherence for both hands (high-frequency) and AEC for the left parietal region (*r* = 0.64, *p* = 0.0444); between motion coherence for PLHCRH (low-frequency) and AEC for the left parietal region (*r* = 0.64, *p* = 0.0412); and between motion coherence for PLHCRH (high-frequency) and AEC for the left parietal region (*r* = 0.76, *p* = 0.0018) (Figure 5).

**Figure 5.**
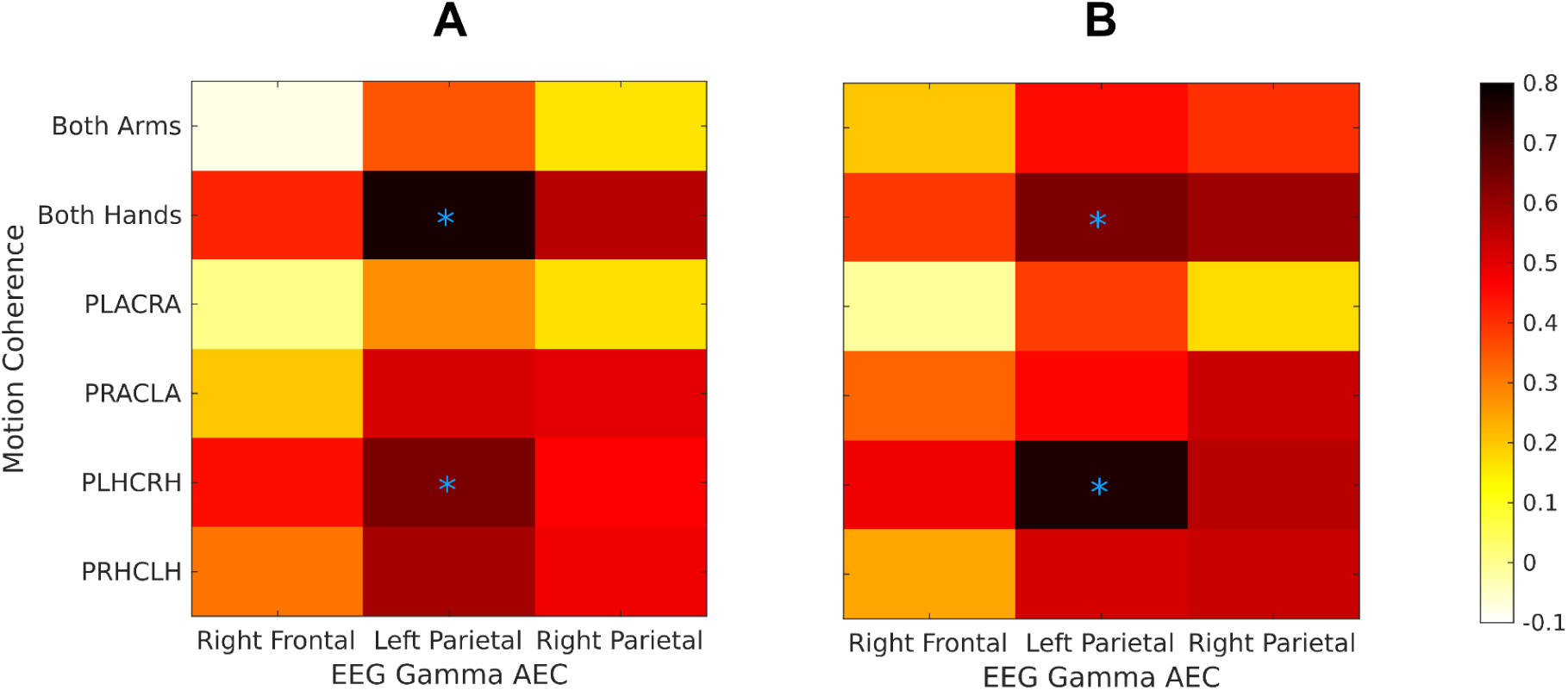
Correlation between measures of low-frequency (A) and high-frequency (B) motion coherence and EEG Gamma AEC in the first mirror game. Asterisks mark significant correlations.

### Correlating motion- and neural synchronization with the Total Test Score

Pearson’s correlation coefficient was calculated between measures of motion synchronization and the Total Test Score and between measures of neural synchronization and the Total Test Score. A significant correlation was found between motion coherence for both hands in the first mirror game in the high-frequency range and the Total Test Score (*r* = 0.58, *p* = 0.0439) and between motion coherence for PRHCLH in the second mirror game in the high-frequency range and the Total Test Score (*r* = 0.63, *p* = 0.0191) (Figure 6).

**Figure 6.**
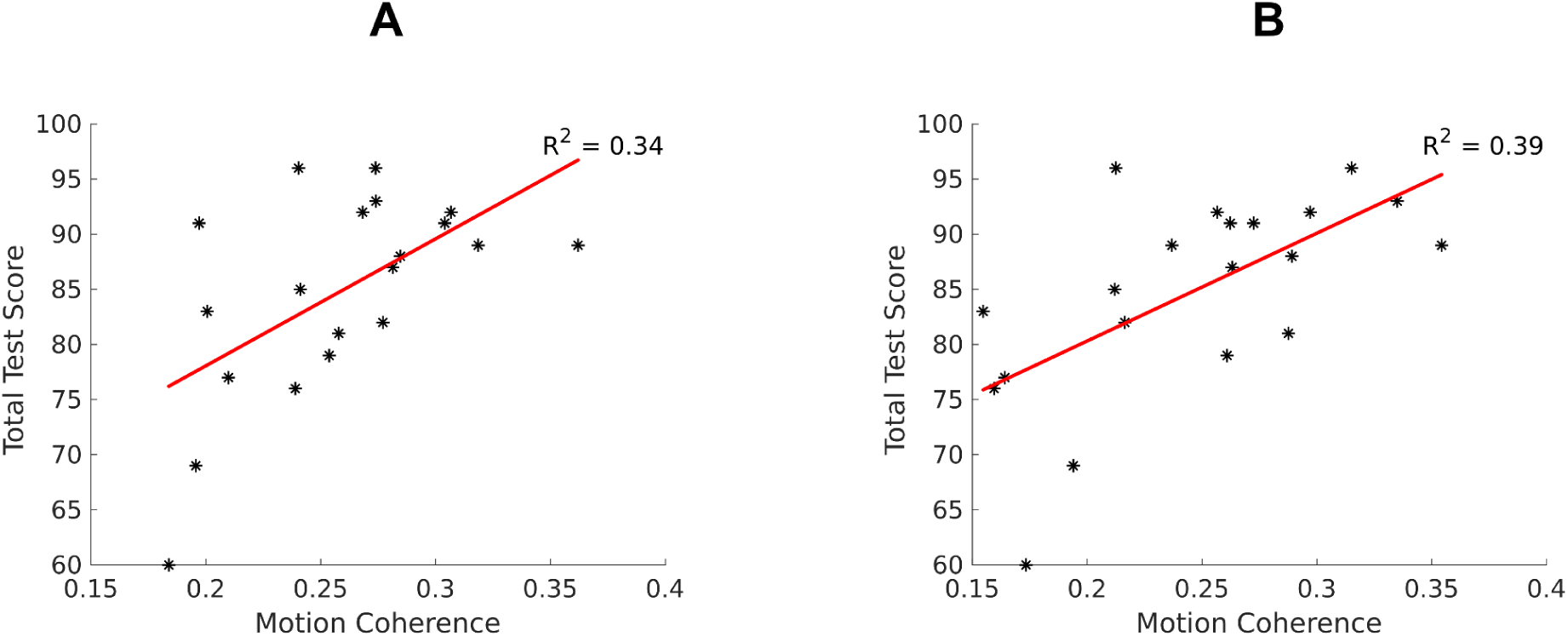
Regression lines of the correlation between motion coherence for both hands in the first mirror game in the high-frequency range and the Total Test Score (A), and between motion coherence for PRHCLH in the second mirror game in the high-frequency range and the Total Test Score (B).

### Correlating motion- and neural synchronization with the times of the labyrinth games

Pearson’s correlation coefficient was calculated between measures of motion synchronization and the times needed to complete the first and the second labyrinth game and between measures of neural synchronization and the times needed to complete the first and the second labyrinth game. Significant negative correlation was found between the time needed to complete the first labyrinth game and four measures of motion synchrony in the first mirror game in the high-frequency range: both arms (*r* = -0.72, *p* = 0.0046), both hands (*r* = -0.75, *p* = 0.0019), PLACRA (*r* = -0.71, *p* = 0.0060), PLHCRH (*r* = -0.62, *p* = 0.0450). Moreover, a significant negative correlation was found between the time needed to complete the second labyrinth game and PLACRA in the first mirror game in the high-frequency range (*r* = -0.62, *p* = 0.0423).

### Comparing the times of the labyrinth games and their correlation with the Total Test Score

The times needed to complete the first and the second labyrinth game were compared by a paired-sample t-test. Mother-child pairs needed significantly less time to solve the labyrinth game in the second repetition compared to the first repetition (*t*(19) = 7.26, *p* < 1e-6, *d* = 1.62). Both times showed a significant correlation with the Total Test Score with a stronger negative correlation for the first labyrinth game (*r* = -0.66, *p* = 0.0029) than for the second labyrinth game (*r* = -0.52, *p* = 0.0357) (Figure 7).

**Figure 7.**
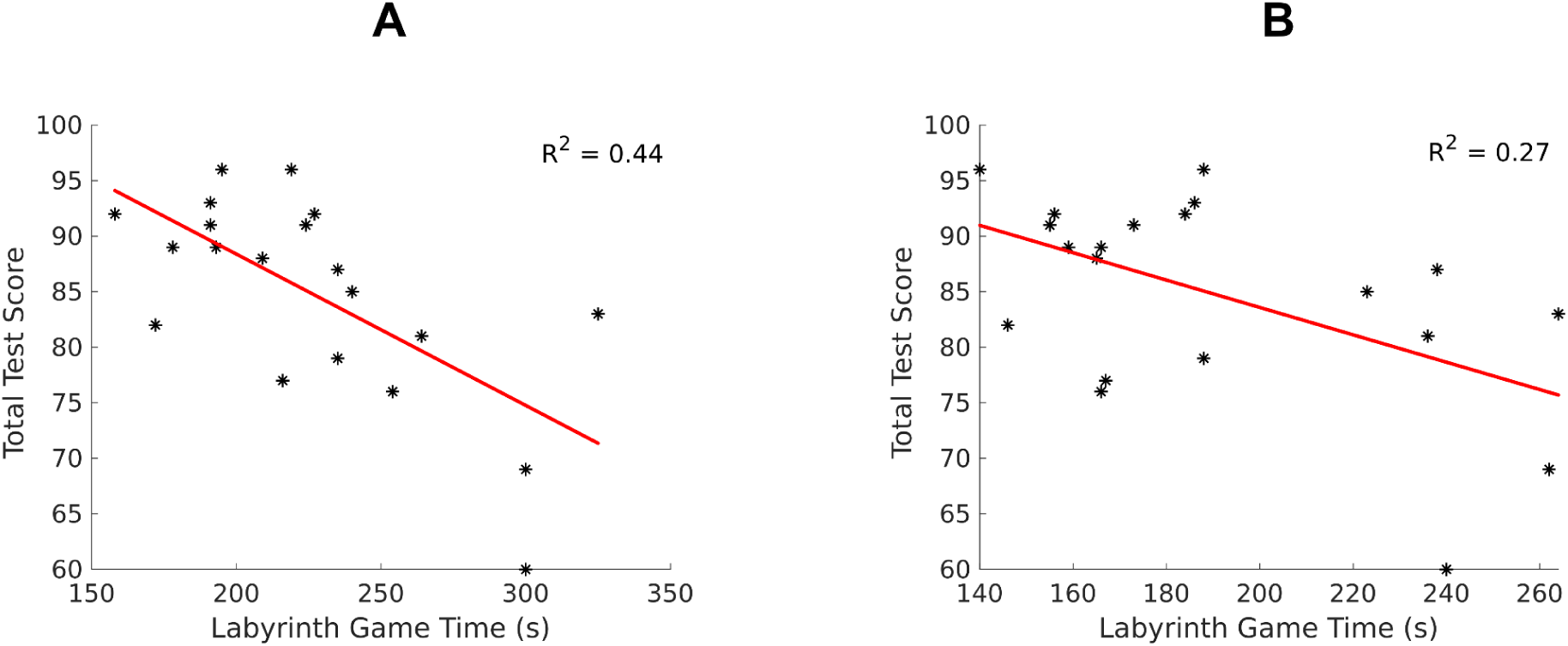
Regression lines of the correlation between the time needed to complete the first labyrinth game and the Total Test Score (A) and between the time needed to complete the second labyrinth game and the Total Test Score (B).

## Discussion

This study investigated brain-to-brain and motor synchronization between mothers and their preschool children during joint motor coordination tasks. Using dual-EEG scanning and upper body motion tracking, we observed significant synchrony in motor actions and neural activity, specifically within the gamma band oscillations. The alignment of motor and neural synchrony within the mother-child dyads evolved between the first and second repetitions of the task, with a strong correlation between motor synchronization and neural coupling. Further analysis revealed that motor synchronization but not neural synchrony was significantly related to the child’s motor development reflected in the MABC total test score. These findings suggest that gamma band brain-to-brain coupling reflects joint motor coordination and mutual adaptation influenced by the motor skills of both the child and the mother. Below, we discuss these results in detail.

### Interpersonal movement synchrony in mother-child interaction

In the present study we found evidence for synchronized movement in mother-child dyads, which is in line with results reported by Cuadros et al (2019) and Feldman et al (2011) who found increased IMS between mother-infant dyads and Bernieri et al (1988) who found the same increase in synchronous activity between mothers and unknown infants. This study extends the findings of previous works by showing that IMS is affected by motor development.

In the mirror game, all six marker groups showed IMS in both first and second repetitions and games and low and high-frequency bands. In contrast, in the labyrinth game, only the high-frequency band showed significantly higher coherence. The difference between the two types of task was that the labyrinth game allowed a lower degree of freedom for the mothers and their child movement while the mirroring game had a higher variance in terms of movements.

Our study was designed to test the hypothesis that the maturity of a child’s movement development correlates with higher levels of motor synchronization, especially in tasks that require imitation. The preschool age group was selected for its relevance in studying motor development, as body schemes that allow for complex imitation are already present but still immature at this stage. We hypothesized that children with more mature body representations would demonstrate faster learning and higher levels of synchrony during joint tasks, particularly in imitation tasks like the mirror game (Jones, 2012; Bremner, 2016). Our findings support this hypothesis, showing that children with more advanced motor skills exhibited stronger movement synchronization and faster task execution across repetitions.

In both the mirror and labyrinth games, motor synchronization was found to correlate with the child’s motor development, with stronger synchrony observed in children who scored higher on motor skills assessments. This suggests that children with more refined body representations can better coordinate their movements with their mothers, leading to higher overall synchrony. The mirror game, which involved more imitation-based tasks with a higher degree of movement variability, showed consistent motor synchrony across both low- and high-frequency bands, further supporting the idea that imitation tasks elicit greater synchronization in children with more advanced motor skills.

This alignment between motor synchrony and motor skill development highlights the importance of motor learning during early childhood. As expected, repetition of the tasks led to faster performance, which was closely linked to the child’s motor abilities. These results align with our hypothesis that children with more mature motor control would not only synchronize better with their mothers but also learn and improve more quickly. This supports the idea that body representations, critical for motor learning and imitation, become more refined with practice, leading to enhanced coordination in joint tasks.

Additionally, this study’s functional relevance of movement synchronization emphasizes its role in social interactions and motor development. As motor synchronization plays a crucial part in social bonding and prosocial behavior (Cirelli et al., 2014), the stronger synchronization seen in children with more developed motor skills suggests a deeper integration of motor and social processes. The findings confirm that synchronization between mother and child reflects motor ability and an indicator of mutual adaptation and shared goals, highlighting the complex interplay between motor learning, social interaction, and emotional connectedness during early development.

Our results align with a growing body of evidence that demonstrates the positive effects of synchronization on learning, particularly through the modulation of attention and the sense of social togetherness (Bevilacqua et al., 2019; Rolf et al., 2009). Studies have consistently shown a positive relationship between social synchrony and enhanced learning performance, suggesting that synchronized activities foster shared attention and cooperation, which drive better outcomes. This is particularly relevant for our study, as the strong correlation we observed between motor synchronization and a child’s motor development supports the idea that synchrony plays a role in facilitating learning, especially in joint motor tasks. However, the relationship between synchrony and performance is not always straightforward. For instance, Vink et al. (2017) found a negative correlation between interpersonal synchrony and performance during a motor task in schoolchildren. This highlights that the nature of the task and the developmental stage of the participants can influence how synchrony impacts learning.

Interestingly, recent research by Pan et al. (2021) showed that transcranial direct current stimulation (tDCS)-induced movement synchrony improved learning performance, further emphasizing the potential role of synchrony in enhancing motor learning. Given that social synchrony has already been shown to increase learning in infants (Rolf et al., 2009), it would be worthwhile to explore whether movement synchrony similarly boosts learning performance in preschool-aged children, a critical period when voluntary movement synchronization is beginning to emerge (Meltzoff & Marshall, 2018; Jones, 2012). Our findings suggest that children with more advanced motor skills demonstrating greater movement synchronization with their mothers may experience enhanced learning and motor development through these coordinated interactions.

The improvement in task execution with repetition, which was correlated with the child’s motor skills, further supports the idea that synchrony aids motor learning. As children refine their body schemes and motor representations, their ability to synchronize movements with others improves, leading to better performance in joint tasks. This aligns with motor learning theories that emphasize the role of repeated practice in enhancing performance (Wolpert et al., 2001). In our study, the higher synchrony observed in children with more developed motor skills may indicate that movement synchrony is not only a byproduct of motor learning but also a facilitator of it, helping children refine their movements through interaction with a more skilled partner, such as their mother.

### Brain oscillations that support joint coordination in mother-child interactions

The present study demonstrates that brain oscillations support inter-brain synchrony during social interactions. We found that parent-child brain synchronization supports joint coordination of the motor task. Specifically, in both tasks, mother-child brain activity was synchronized exclusively in gamma band oscillatory activity. Interbrain synchrony is strongly associated with behavioral synchrony, social gaze, and positive affect, highlighting the importance of social connectedness in two-brain coordination (Kinreich et al., 2017). The degree of familiarity and attachment between individuals influences the efficiency of neural synchronization, with romantic couples typically demonstrating higher levels of brain-to-brain synchrony compared to strangers (Kinreich et al., 2017; Djalovski et al., 2020; Müller & Lindenberger, 2014). These findings suggest that human attachments provide a framework for the efficient, resource-sensitive performance of social goals through coordinated brain activity. We found gamma band brain-to-brain coupling during motor coordination tasks between mothers and preschool children, likely because this frequency range is closely associated with social and behavioral synchrony in naturalistic interactions. Previous research has shown that during social interactions, neural synchrony often emerges in gamma rhythms, particularly in temporal-parietal regions. Studies on romantic couples have demonstrated that gamma band synchronization correlates with behavioral synchrony, social gaze, and positive affect, highlighting the role of social connectedness in coordinating two brains (Kinreich et al., 2017). Similarly, the strong emotional and social bond between a mother and child may activate gamma rhythms, particularly when engaging in joint motor tasks that require attention, coordination, and mutual adaptation. This suggests that the gamma band synchrony observed in our study may reflect the intense interpersonal engagement required during motor coordination.

Moreover, research on parent-child dyads emphasizes the role of neural synchrony in early childhood development and social interactions. While brain-to-brain synchrony in adults typically occurs in the alpha-mu band during social coordination tasks (Dumas et al., 2012), younger children may rely more on gamma oscillations due to ongoing cortical development. Neural synchrony, including gamma rhythms, plays a critical role in the maturation of cortical networks during childhood, supporting joint attention, emotion regulation, and social bonding (Uhlhaas et al., 2009). The dynamic and interactive nature of mother-child relationships may elicit gamma band coupling as it reflects the need for real-time, adaptive coordination during motor tasks. This synchronization in the gamma band may serve as a developmental foundation for more complex forms of social interaction, such as joint action and shared intentionality, which evolve as children grow older.

The observed gamma band interbrain synchronization is localized to the frontal (right hemisphere) bilateral parietal cortices in the first mirror game. In the first labyrinth game, the bilateral central and right parietal regions synchronize between the mother and child’s gamma band brain activity. Thus, we suggest that the observed brain-to-brain synchrony likely reflects the nature of the social interaction, with specific brain areas such as temporal-parietal and sensorimotor regions becoming involved depending on the task’s demands, as previous studies have shown similar patterns during coordinated activities (Kinreich et al., 2017; Djalovski et al., 2020).

Studies have shown that neural synchronization emerges in various frequency bands, particularly in the alpha-mu (Dumas et al., 2010), beta, and gamma rhythms (Kinreich et al., 2017; Djalovski et al., 2020). We did not find alpha band interbrain synchrony in the present study. Alpha band activity (typically in the 8-12 Hz range) has been extensively studied in relation to cognitive processes like attention, social cognition, and sensorimotor integration. Alpha band synchrony often reflects shared attention, joint action, or mutual engagement, making it a frequently employed index in studies of interpersonal brain-to-brain coupling. It is possible that the finding of gamma band interpersonal synchrony, along with the lack of alpha band synchrony, is related to the fact that the study involved preschool-aged children interacting with their mothers. Several factors could account for this. First, neural oscillations, including those in the alpha band, undergo significant development during early childhood.

Preschool-aged children may not exhibit alpha band synchrony like adults, as their neural systems are still maturing. In contrast, gamma band activity, associated with attention, perception, and cognitive processing, may be more prominent in young children during interactive tasks. Additionally, the specific motor tasks investigated might have required a higher level of joint attention, active coordination, and rapid processing—cognitive functions more strongly linked to gamma band activity. While alpha band synchrony is often observed in adults during sustained attention and social interaction, the dynamic nature of preschool children’s interactions may engage gamma oscillations more intensively. Lastly, the emotional and social bonding between mothers and their children may evoke neural processes more associated with gamma band synchrony, reflecting intense joint engagement and coordination. In contrast, the typical alpha band synchrony seen in adult dyads during cooperative tasks may be less prominent in the highly interactive and rapidly changing context of mother-child interactions. Therefore, the absence of alpha band synchrony could result from both the children’s developmental stage and the specific dynamics of mother-child interactions in the study.

Neural synchrony within the mother-child dyad’s brain activities is generally higher at the beginning of the task and declines with the repetition of motor tasks, regardless of which type of task we measured. An alternative interpretation of the results could suggest that the higher neural synchrony observed at the beginning of the task reflects initial mutual engagement or shared cognitive effort as the mother and child coordinate their actions. As the task progresses and becomes more familiar through repetition, the need for close neural alignment decreases, possibly due to the dyad’s increasing individual autonomy and task proficiency. This decline in synchrony may indicate a shift from a highly interactive, joint problem-solving mode to more independent task execution as both participants internalize the motor sequences.

We found that motor synchronization during mother-child motor tasks is correlated with the child’s trait motor skills measured with a developmental assessment battery. There was a strong positive relation between brain-to-brain coupling and motor synchrony during the task. However, the child’s motor skills did not correlate with the gamma band brain-to-brain coupling. One alternative explanation for the lack of correlation between the child’s motor skills and gamma band brain-to-brain coupling could be that neural synchrony reflects interactive processes that emerge specifically during the social interaction rather than directly tied to the child’s motor abilities. Brain-to-brain coupling may be more sensitive to the dynamic coordination between the mother and child, which involves mutual adaptation, shared goals, and emotional understanding beyond the child’s motor skills. Additionally, the mother’s influence, including her experience and ability to adjust to the child’s movements, may play a crucial role in fostering this neural synchrony, indicating that brain-to-brain coupling is shaped by joint interaction rather than just the child’s motor competence. This suggests that neural synchrony could be more reflective of the relational and collaborative nature of the task rather than the static traits of either individual.

These findings suggest that movement synchronization during motor tasks may enhance both motor development and learning in preschool-aged children, particularly as they begin to master voluntary coordination. Gamma band brain-to-brain coupling during these tasks may reflect the underlying neural processes that support this synchronization, pointing to the functional relevance of movement synchrony in facilitating joint attention, cooperation, and learning in early childhood. Further research could explore how these synchronized interactions contribute to long-term developmental outcomes, particularly as voluntary motor synchronization matures throughout early childhood.

### Limitations and Future Direction

While the study provides insights into the relationship between motor-neural synchronization and learning, it has limitations. The sample size of 20 mother-child dyads, though sufficient for detecting significant effects, limits the generalizability of the findings. Larger-scale studies are needed to confirm these results across different populations and to explore potential gender differences or the influence of individual characteristics (e.g., temperament, personality). Moreover, while the study focused on mother-child interactions, future research could examine how synchrony dynamics differ across caregiver-child dyads (e.g., fathers, grandparents).

A key limitation of analysis methods in the current study is the need for more insight into the temporal dynamics, short-term structure, and synchronization evolution within the tasks. The current study confirms a measurable and meaningful interaction between behavioral and neural data. However, the intricacies of these interactions will only come to light using measures that can accurately quantify synchrony over shorter periods and a wider range of frequencies, such as cross-wavelet and wavelet coherence measures. (e.g. Bigand et al., 2024) Another advantage of such methods is that motion and EEG data can be seamlessly combined in the analysis (Issartel et al., 2015).

A further limitation is focusing on only two tasks (the Mirror Game and the Labyrinth Game). While these tasks provide a valuable contrast between different levels of task structure, expanding the range of tasks to include more complex or naturalistic social interactions would offer a more comprehensive understanding of how synchrony evolves across different contexts. Longitudinal studies would also be valuable in tracking how synchrony develops over time and its long-term impact on motor and cognitive development.

## Conclusions

The study’s findings underscore the critical role of both motor and neural synchronization in early childhood development, particularly in the context of motor learning and parent-child interaction. Synchrony not only facilitates joint actions but also serves as a marker of developmental progress, with implications for early intervention and education. By understanding the dynamics of synchrony, we can gain valuable insights into the neural and behavioral foundations of learning and social interaction, ultimately contributing to more effective strategies for supporting early childhood development.

## Data Statement

The data in this study contain sensitive information (audio and video recordings) about participants, some of whom are members of a vulnerable group (children). Therefore, the raw data will not be made publicly available. Anonymized data are available from the corresponding author upon request.

## Acknowledgment

The authors are grateful to the RCNS staff (Emese Várkonyi, Györgyi Balla, Eszter Rozgonyiné Lányi) who recorded the data and, finally, to all participants and their families.

## Ethics

The study was conducted according to the Declaration of Helsinki and all applicable national laws, and it was approved by the relevant ethics committee: the United Ethical Review Committee for Research in Psychology (EPKEB) in Hungary (Ref. no: 2022-113).

