## Supplementary tables for "Interpersonal Synchronization in Mother-Child Dyads: Neural and Motor Coupling as a Mechanism for Motor Learning and Development in Preschoolers"

**Supplementary materials**

Table A1: Results of two-sample t-tests comparing the coherence of real and pseudo pairs in the low-frequency band (0.1 - 1 Hz) in the first mirror game. Significance values are reported with Bonferroni-correction against multiple comparisons.

| Marker group | *t* | *df* | *p* | *d* |
|---|---|---|---|---|
| Both arms | 4.62 | 398 | 0.00003 | 1.06 |
| Both hands | 8.34 | 398 | <1e-6 | 1.91 |
| PLACRA | 6.45 | 398 | <1e-6 | 1.48 |
| PRACLA | 9.04 | 398 | <1e-6 | 2.07 |
| PLHCRH | 12.13 | 398 | <1e-6 | 2.78 |
| PRHCLH | 12.50 | 398 | <1e-6 | 2.87 |

Table A2: Results of two-sample t-tests comparing the coherence of real and pseudo pairs in the high-frequency band (1 - 4 Hz) in the first mirror game. Significance values are reported with Bonferroni-correction against multiple comparisons.

| Marker group | *t* | *df* | *p* | *d* |
|---|---|---|---|---|
| Both arms | 4.41 | 398 | 0.00008 | 1.01 |
| Both hands | 9.24 | 398 | <1e-6 | 2.12 |
| PLACRA | 7.06 | 398 | <1e-6 | 1.62 |
| PRACLA | 6.75 | 398 | <1e-6 | 1.55 |
| PLHCRH | 11.27 | 398 | <1e-6 | 2.59 |
| PRHCLH | 11.28 | 398 | <1e-6 | 2.59 |

Table A3: Results of two-sample t-tests comparing the coherence of real and pseudo pairs in the low-frequency band (0.1 - 1 Hz) in the second mirror game. Significance values are reported with Bonferroni-correction against multiple comparisons.

| Marker group | *t* | *df* | *p* | *d* |
|---|---|---|---|---|
| Both arms | 4.88 | 398 | 0.000009 | 1.12 |
| Both hands | 7.69 | 398 | <1e-6 | 1.76 |
| PLACRA | 5.54 | 398 | <1e-6 | 1.27 |
| PRACLA | 7.48 | 398 | <1e-6 | 1.72 |
| PLHCRH | 9.42 | 398 | <1e-6 | 2.16 |
| PRHCLH | 12.87 | 398 | <1e-6 | 2.95 |

Table A4: Results of two-sample t-tests comparing the coherence of real and pseudo pairs in the high-frequency band (1 - 4 Hz) in the second mirror game. Significance values are reported with Bonferroni-correction against multiple comparisons.

| Marker group | *t* | *df* | *p* | *d* |
|---|---|---|---|---|
| Both arms | 4.53 | 398 | 0.00005 | 1.04 |
| Both hands | 5.47 | 398 | <1e-6 | 1.25 |
| PLACRA | 7.13 | 398 | <1e-6 | 1.64 |
| PRACLA | 6.48 | 398 | <1e-6 | 1.49 |
| PLHCRH | 8.72 | 398 | <1e-6 | 2.00 |
| PRHCLH | 8.74 | 398 | <1e-6 | 2.01 |

Table A5: Results of two-sample t-tests comparing the coherence of real and pseudo pairs in the low-frequency band (0.1 - 1 Hz) in the first labyrinth game. Significance values are reported with Bonferroni-correction against multiple comparisons.

| Marker group | *t* | *df* | *p* | *d* |
|---|---|---|---|---|
| Both arms | 0.93 | 398 | >0.5 | 0.21 |
| Both hands | 2.15 | 398 | 0.19 | 0.49 |
| PLACRA | 1.80 | 398 | 0.43 | 0.41 |
| PRACLA | -0.51 | 398 | >0.5 | -0.12 |

| | | | | |
|---|---|---|---|---|
| PLHCRH | 1.28 | 398 | >0.5 | 0.29 |
| PRHCLH | 0.35 | 398 | >0.5 | 0.08 |

Table A6: Results of two-sample t-tests comparing the coherence of real and pseudo pairs in the high-frequency band (1 - 4 Hz) in the first labyrinth game. Significance values are reported with Bonferroni-correction against multiple comparisons.

| Marker group | *t* | *df* | *p* | *d* |
|---|---|---|---|---|
| Both arms | 3.53 | 398 | 0.0028 | 0.81 |
| Both hands | 4.68 | 398 | 0.00002 | 1.07 |
| PLACRA | 5.90 | 398 | <1e-6 | 1.35 |
| PRACLA | 2.63 | 398 | 0.0533 | 0.60 |
| PLHCRH | 6.71 | 398 | <1e-6 | 1.54 |
| PRHCLH | 4.46 | 398 | 0.00006 | 1.02 |

Table A7: Results of two-sample t-tests comparing the coherence of real and pseudo pairs in the low-frequency band (0.1 - 1 Hz) in the second labyrinth game. Significance values are reported with Bonferroni-correction against multiple comparisons.

| Marker group | *t* | *df* | *p* | *d* |
|---|---|---|---|---|
| Both arms | 0.92 | 398 | >0.5 | 0.21 |
| Both hands | 1.24 | 398 | >0.5 | 0.29 |
| PLACRA | 0.10 | 398 | >0.5 | 0.02 |
| PRACLA | 1.47 | 398 | >0.5 | 0.34 |
| PLHCRH | 0.49 | 398 | >0.5 | 0.11 |
| PRHCLH | 1.45 | 398 | >0.5 | 0.33 |

Table A8: Results of two-sample t-tests comparing the coherence of real and pseudo pairs in the high-frequency band (1 - 4 Hz) in the second labyrinth game. Significance values are reported with Bonferroni-correction against multiple comparisons.

| Marker group | *t* | *df* | *p* | *d* |
|---|---|---|---|---|
| Both arms | 3.06 | 398 | 0.014 | 0.70 |
| Both hands | 3.38 | 398 | 0.0047 | 0.78 |
| PLACRA | 1.92 | 398 | 0.33 | 0.44 |
| PRACLA | 3.45 | 398 | 0.0037 | 0.79 |
| PLHCRH | 1.98 | 398 | 0.29 | 0.45 |
| PRHCLH | 3.82 | 398 | 0.0009 | 0.88 |
